# Ultradian Rhythms in Heart Rate Variability and Distal Body Temperature Anticipate the Luteinizing Hormone Surge Onset

**DOI:** 10.1101/2020.07.15.205450

**Authors:** Azure D. Grant, Mark Newman, Lance J. Kriegsfeld

## Abstract

The human menstrual cycle is characterized by predictable patterns of physiological change across timescales, yet non-invasive anticipation of key events is not yet possible at individual resolution. Although patterns of reproductive hormones across the menstrual cycle have been well characterized, monitoring these measures repeatedly to anticipate the preovulatory luteinizing hormone (LH) surge is not practical for fertility awareness. In the present study, we explored whether non-invasive and high frequency measures of distal body temperature (DBT), sleeping heart rate (HR), sleeping heart rate variability (HRV), and sleep timing could be used to anticipate the preovulatory LH surge in women. To test this possibility, we used signal processing to examine these measures across the menstrual cycle. Cycles were examined from both pre- (n=45 cycles) and perimenopausal (n=10 cycles) women using days of supra-surge threshold LH and dates of menstruation for all cycles. For a subset of cycles, urinary estradiol and progesterone metabolites were measured daily around the time of the LH surge. Wavelet analysis revealed a consistent inflection point of ultradian rhythm (2-5 h) power of DBT and HRV that enabled anticipation of the LH surge at least 2 days prior to its onset in 100% of individuals. In contrast, the power of ultradian rhythms in heart rate, circadian rhythms in body temperature, and metrics of sleep duration and sleep timing were not predictive of the LH surge. Together, the present findings reveal fluctuations in distal body temperature and heart rate variability that consistently anticipate the LH surge and may aid in fertility awareness.

**Key Points:**

- Ultradian (2-5 h) rhythm power of distal body temperature and heart rate variability (RMSSD) exhibits a stereotyped inflection point and peak in the days leading up to the LH surge in premenopausal women.
- Circadian rhythms of distal body temperature and single time-point/day metrics do not permit anticipation of the LH surge.
- Measurement of continuous metabolic and autonomic outputs, enabling assessment of ultradian rhythms, may be of value to the fertility awareness method.

## Introduction

The fertility-awareness-method (FAM), a set of practices used to estimate the fertile and infertile days of the ovulatory cycle, is challenging to implement and to study, and existing studies of its effectiveness are inconclusive (Grimes *et al*., 2005). However, an observation-based method of family planning or contraception has several potential benefits, including a lack of hormonal disruption, personalization and relatively low cost. One challenge inherent to current FAM practices is the reliance on historical basal body temperature and symptom trends (e.g., breast tenderness, libido, cervical mucous) that can vary substantially by individual, and within-individual from cycle-to-cycle (Shilaih *et al*., 2017), and that provide predominantly retrospective information. The challenges of FAM have led the majority of those seeking to avoid pregnancy to adopt another form of contraception. Unfortunately, the most widely used form of contraception, female hormonal contraception, has short and long term risks for many users, including but not limited to increased breast cancer rate (Mørch *et al*., 2017; Marsden, 2017), luteal phase deficiency (Gnoth *et al*., 2002), dysmenorrhea (Delbarge *et al*., 2002; Gnoth *et al*., 2002), altered cognition (Pletzer & Kerschbaum, 2014; Bradshaw *et al*., 2020) and depressed mood (Skovlund *et al*., 2016, 2018). These risks, combined with increasing recognition that physiological systems vary in a structured manner across the menstrual cycle (Brar *et al*., 2015; Smarr *et al*., 2016, 2017; Grant *et al*., 2018), provide the impetus to develop FAM approaches that employ high-temporal-resolution, non-invasive measures of physiology.

The menstrual cycle is a continuous, rhythmic succession of endocrine, ovarian and uterine events. Briefly, the cycle begins with onset of menstruation, followed by rising levels of estradiol, follicular maturation and proliferation of the uterine lining (Baerwald *et al*., 2012; Ferreira & Motta, 2018). Ovulation, which is triggered by numerous factors including estradiol, a surge of LH, the presence of a mature Graafian follicle, and likely time of day (Simonneaux *et al*., 2017), frequently occurs between 1/2 and 3/4 of the way through the cycle in humans (Robker *et al*., 2018). Other physiological systems, including metabolism (Yeung *et al*., 2010; Zhang *et al*., 2020) and autonomic balance (Tada *et al*., 2017) fluctuate with the menstrual cycle. An individual is mostly likely to become pregnant during the time leading up to, and shortly past, the ovulation event, making identification of this period central for the successful use of the FAM. Although high-frequency hormone measurements (e.g., daily estradiol from blood or urine) and ultrasound can provide information on when an LH surge and subsequent ovulation are likely to occur, such measurements are both laborious and expensive, limiting their widespread utility. Furthermore, at home tests available for measuring supra-threshold LH concentrations provide retrospective rather than prospective information for this event. Ideally, new methods of fertility awareness would accurately indicate the approaching peri-ovulatory period via relatively inexpensive and non-invasive means (Stanford, 2015). This study aimed to develop such a preliminary indicator for future, larger scale investigation.

The premise of the present investigation is that the presence of structured changes to peripheral biological rhythms across the menstrual cycle may allow for anticipation of the LH surge. Such a finding would further support the notion that the state of one system (e.g., reproductive) can be inferred via measurements of another (e.g., autonomic or metabolic) (Shannahoff-Khalsa *et al*., 1996; Grant *et al*., 2018; Goh *et al*., 2019). Perhaps the most consistent biological rhythmic changes across the menstrual cycle occur at the 1-4 h (ultradian) timescale (Brandenberger *et al*., 1987; Shannahoff-Khalsa *et al*., 1996, 1997; Grant *et al*., 2018; Zavala *et al*., 2019). Most elements of the hypothalamic-pituitary-ovarian axis, including gonadotropin releasing hormone (GnRH) (Clarke *et al*., 1987; Moenter *et al*., 1991; Gore *et al*., 2004), luteinizing hormone (LH) (Backstrom *et al*., 1982; Vugt *et al*., 1984; Rossmanith *et al*., 1990), FSH (Yen *et al*., 1972; Genazzani *et al*., 1993; Booth Jr *et al*., 1996; Pincus *et al*., 1998), estradiol (Backstrom *et al*., 1982; Licinio *et al*., 1998), progesterone (Backstrom *et al*., 1982; Filicori *et al*., 1984; Veldhuis *et al*., 1988; Soules *et al*., 1989; Rossmanith *et al*., 1990; Genazzani *et al*., 1991) and testosterone (Nóbrega *et al*., 2009) show ultradian rhythms (URs) that are coordinated with menstrual phase (Grant *et al*., 2018). Across species, patterns of these neuropeptides and hormones exhibit an increase in ultradian frequency and inter-hormone coupling strength leading up to ovulation (Rossmanith *et al*., 1990; Moenter *et al*., 1991) and a decrease in ultradian frequency and stability in the luteal phase (Backstrom *et al*., 1982; Healy *et al*., 1984; Filicori *et al*., 1984; Vugt *et al*., 1984; Rossmanith *et al*., 1990; Genazzani *et al*., 1991; Moenter *et al*., 1991; Licinio *et al*., 1998). Additionally, peripheral measures of distal body temperature (DBT) and heart rate variability (HRV) reflect the activity of reproductive (Leicht *et al*., 2003; Chen *et al*., 2008; Shechter *et al*., 2011), autonomic (Shannahoff-Khalsa *et al*., 1996; Stein *et al*., 2006*b*, 2006*a*; Visrutha *et al*., 2012; de Zambotti *et al*., 2013, 2015; Tada *et al*., 2017; Charkoudian *et al*., 2017) and metabolic systems (Buxton & Atkinson, 1948; Shannahoff-Khalsa *et al*., 1996; Fredholm *et al*., 2011; Stuckey *et al*., 2015) and show both URs and menstrual rhythms (Shechter *et al*., 2011). These peripheral and endocrine measures are proposed to operate as coupled oscillators at the ultradian time scale. Assessment of these measures could therefore, potentially enable endocrine status assessment via timeseries analysis of peripheral measures (Shannahoff-Khalsa *et al*., 1996; Grant *et al*., 2018; Goh *et al*., 2019).

Recent work in animals supports the idea that non-reproductive measures can be used to anticipate reproductive status. In rodents, the power of core body temperature URs exhibits a trough on the day of ovulation (Smarr *et al*., 2016, 2017). The translational capability of this method is supported by the association of gross timescale changes in DBT, HR and HRV by menstrual phase (Buxton & Atkinson, 1948; Sato *et al*., 1995; Leicht *et al*., 2003; Yildirir *et al*., 2006; Shechter *et al*., 2011; Uckuyu *et al*., 2013; Kräuchi *et al*., 2014; Brar *et al*., 2015; Uchida *et al*., 2019; Zhang *et al*., 2020). However, it is unknown if patterns of rhythmic change from continuous measures leading up to the LH surge associate with human reproductive status. Although the specific factors responsible for the changes in frequency of reproductive URs across non-human mammalian ovulatory cycles are not well understood, their consistency of change across species of widely varying cycle length suggests a concerted role in menstrual cycle function (Grant *et al*., 2018). Finally, the structure of some circadian rhythms (∼24 h; CRs) is altered in the luteal phase, with estradiol acrophase advancing, and REM sleep exhibiting a modest decrease; however, structured changes are not generally observed during the peri-ovulatory period (Baker & Driver, 2007 p.200). As both URs and CRs are tightly regulated across systems, monitoring their structure may enable more accurate assessment of reproductive state than infrequently collected data (e.g., 1 time point per day) (Bauman, 1981; Quagliarello & Arny, 1986; Zavala *et al*., 2019).

Wearable devices offer unprecedented ease of collecting the continuous, longitudinal data needed to assess URs and CRs across the menstrual cycle (Gear, n.d.; Hasselberg *et al*., 2013; de Zambotti *et al*., 2017; Epstein *et al*., 2017). To determine if rhythmic structure exhibits reliable changes leading up to the LH surge, we used a wearable device (the Oura© Ring) to monitor distal body temperature (DBT) continuously, sleeping heart rate (HR), sleeping heart rate variability (HRV (root mean square of successive differences; RMSSD)), sleep timing and duration. If endocrine, metabolic and autonomic rhythms are sufficiently coupled at the ultradian and circadian timescales, then coordinated patterns should be observed across measures and potentially across the menstrual cycle. Such patterns would contribute to a growing body of work in “network physiology” (Bashan *et al*., 2012; Bartsch *et al*., 2015), which proposes that changes among endocrine, metabolic and autonomic outputs are coupled under real world conditions. As mentioned above, implicit in this hypothesis is that one could infer the state of one system via measurements of another. Anticipation of female reproductive events is a test of the network physiology framework with potential for rapid translation.

## Materials and Methods

### Ethical Approval

This study and all procedures were approved by the Office for the Protection of Human Subjects at the University of California, Berkeley. All participants gave informed consent.

### Participants and Recruitment

Participants were recruited from the Quantified Self community, a global group of individuals interested in learning through self-measurement (Choe *et al*., 2014; Grant *et al*., 2019). Individuals attended a prospective discussion about the project at the 2018 Quantified Self meeting in Portland, Oregon and contacted the experimenter via email if interested in participating. Prospective participants were contacted to discuss study structure, risks and benefits, and to review the informed consent form. Once informed consent was obtained, participants were instructed to complete an introductory questionnaire with their age, cycling status (regular, irregular, recovering from hormonal/IUD contraceptive use, amenorrhea, perimenopausal, menopausal), and historical LH surge day(s), if known. Contact information was collected for the purposes of communication and delivery of study materials. There were no exclusion criteria, but data from pregnancies (*n* =3) that overlapped with the study were excluded from these analyses. Participants had not taken hormonal contraception within the prior year and did not have any known reproductive medical concerns. There were no age or parity restrictions, consistent with the principles of participatory research (Vayena & Tasioulas, 2013; Grant *et al*., 2019). See **Table 1** for participant characteristics.

### Study Design

Each of the 28 (*n*=20 premenopausal, *n*=5 perimenopausal, *n*=3 premenopausal and became pregnant) participants collected 2 to 3 cycles of data for analysis. For all cycles, the Oura© Ring a distal body temperature (DBT), HRV (RMSSD), Heart Rate (HR) and sleep sensor, was worn continuously on the finger, as previously described (de Zambotti *et al*., 2017; Maijala *et al*., 2019). For all cycles, LH was monitored via urinary test strips (Wondfo Biotech Co., Guangzhou, China) from day 10 (with first day of menstruation considered day 0) until a positive reading was detected, and subsequently until 2 days after LH fell below the limit of detection (see below for details on the *Urinary Hormone Assay, Luteinizing Hormone*). Of the 55 total cycles collected (45=premenopausal, 10=perimenopausal), 20 were paired with daily, morning urine tests for estradiol, α-pregnanediol (αPg) and β-pregnanediol (βPg), the major urinary progesterone metabolites (Precision Analytical, McMinnville, OR). This study was designed using the principles of Participant-Led Research (PLR) (Grant *et al*., 2019; Grant & Wolf, 2019) in which individual participants maintain control of their own data prior to anonymization and came to the project with personal questions that could be answered with the data to be collected. All participants received a copy of their Oura© Ring data.

### Data Collection and Management

HR, HRV (RMSSD), DBT, sleep onset, sleep offset, sleep duration, breathing rate and nightly average temperature deviation (described briefly below) were collected using the Oura© Ring (Oura Inc., San Francisco, CA). The Oura© ring is a small, wireless sensor worn on the finger. By using an LED light source and LED sensor to measure reflection off the skin above the radial artery of the finger, the Oura© ring calculates HR, HRV (RMSSD) and breathing rate. The ring also contains 3 thermistors for detection of DBT. DBT is measured 24 hours a day (binned in 5-minute intervals), whereas HR and HRV (RMSSD) are only measured during sleep (also binned in 5-minute intervals), limiting our analyses of HR and HRV (RMSSD) to the sleeping period. All other metrics are calculated once per night. Briefly, the body temperature deviation for each night is the average nightly temperature between 10:00pm and 8:00am, minus the average temperature of the previous 20 days. Oura© rings were loaned to the group by Oura Inc.

The Oura© ring can be connected to a mobile phone application, Oura, via Bluetooth. At the start of the study, each participant downloaded the Oura application from either the Google Play Store (Google Inc., Mountain View, CA) or the Apple App Store (Apple Inc, Cupertino, CA) to their mobile phones and created an Oura account. Participants were able to view their own data provided by the application throughout the study. Participants were asked to synchronize data from the ring to the application each morning. Uploaded data was automatically transferred via the internet to the study database in the Oura cloud service. In order to access data from the cloud, data were imported into In the Open Humans (Greshake Tzovaras *et al*., 2019) framework, which provides encrypted, password protected data access to researchers, with the participants’ revocable consent. In addition to data collected by the Oura ring, participants uploaded personal spreadsheets that tracked days of menstruation, days of LH tests and results, days of urine collection, and notes (e.g., forgot to wear the Oura ring) to Open Humans. Participants could opt out of the study and remove their data at any time. Data were anonymized by the researchers for analysis. Data, once anonymized at the end of the study, remained in the data set.

### Hormone Assays

For the assessment of E2, αPg and βPg, participants collected daily, first-morning urine samples across a cycle according to manufacturer’s instructions (Precision Analytical, Willamette, OR). Briefly, a standardized piece of filter paper with an attached label was submerged in the urine sample and dried for 24 h. Filter paper was then frozen at ∼ −18°C in participants’ home freezers until analysis. E2, αPg and βPg were analyzed using proprietary in-house assays referred to as Dried Urine Testing for Comprehensive Hormones (DUTCH) on the Agilent 7890/7000B GC–MS/MS (Agilent Technologies, Santa Clara, CA, USA). The equivalent of approximately 600 μl of urine was extracted from the filter paper using acetate buffer. In the first week of the cycle, and from 3 days after LH surge completion until the end of the luteal phase, samples were pooled every 2 days (a third day was pooled at the end of cycles in instances where the total number of remaining days after the surge was odd). Urine samples were extracted and analyzed as previously described, with previously established ranges of hormone concentrations expected in urine by phase of cycle and during menopause (Roos *et al*., 2015; Newman *et al*., 2019). Briefly, creatinine was measured in duplicate using a conventional colorimetric (Jaffe) assay. Conjugated hormones were extracted (C18 solid phase extraction), hydrolyzed by Helix pomatia) and derivatized prior to injection (GC-MS/MS) and analysis. The average inter-assay coefficients of variation were 8% for E2, 12% for αPg, and 13% for βPg. The average intra-assay coefficients of variation were 7% for E2, 12% for αPg and 12% for βPg. Sensitivities of the assays were as follows: E2 and αPg, 0.2 ng/mL; βPg, 10 ng/ mL.

Luteinizing hormone was measured using the commercially available WondFo© (Wondfo Biotech Co., Guangzhou, China) Luteinizing Hormone Urinary Test (Barron *et al*., 2018), a validated at-home urine assay. Briefly, the strip was submerged by participants for 5 seconds in a fresh urine sample and laid horizontally for 5 min before reading. Samples collected for E2, αPg and βPg, were also used for LH testing. Each strip contains a positive control and a “test” line, indicating if LH is present in the urine at, or over a concentration of 25 MIU/mL (Barron *et al*., 2018). Test results were depicted as either + or – (no quantitative information provided) and were recorded in a personal spreadsheet by the participant. A photograph of each test was taken by participants to ensure accurate reading of the results.

### Inclusion and Exclusion Criteria for Collected Cycles

Cycles were included in the premenopausal data set as likely ovulatory by four criteria a) one or more localized days of supra-threshold LH concentration, b) the presence of a rise in E2 (if collected) within typical range prior to or coincident with supra threshold LH, c) a subsequent rise in αPg and βPg (if collected), and d) positive values of DBT deviation, as previously described (Maijala *et al*., 2019), within 2 days of surge onset until the end of the cycle (See **Supplemental Figure 1**). Cycles without E2, αPg and βPg data were included by meeting criteria a and d only. Cycles with missing data within sixteen days of the of the LH surge (defined as no HR/HRV/DBT data for a given night) were omitted in order to avoid erroneous estimation of rhythmic power (see *Data Analysis* below). Cycles were defined as “perimenopausal” by the presence of positive LH measured at least every other day across the cycle and age >45 years. Four such cycles were paired with daily urinary hormone analysis for E2, αPg and βP, as described above.

**Figure 1.**
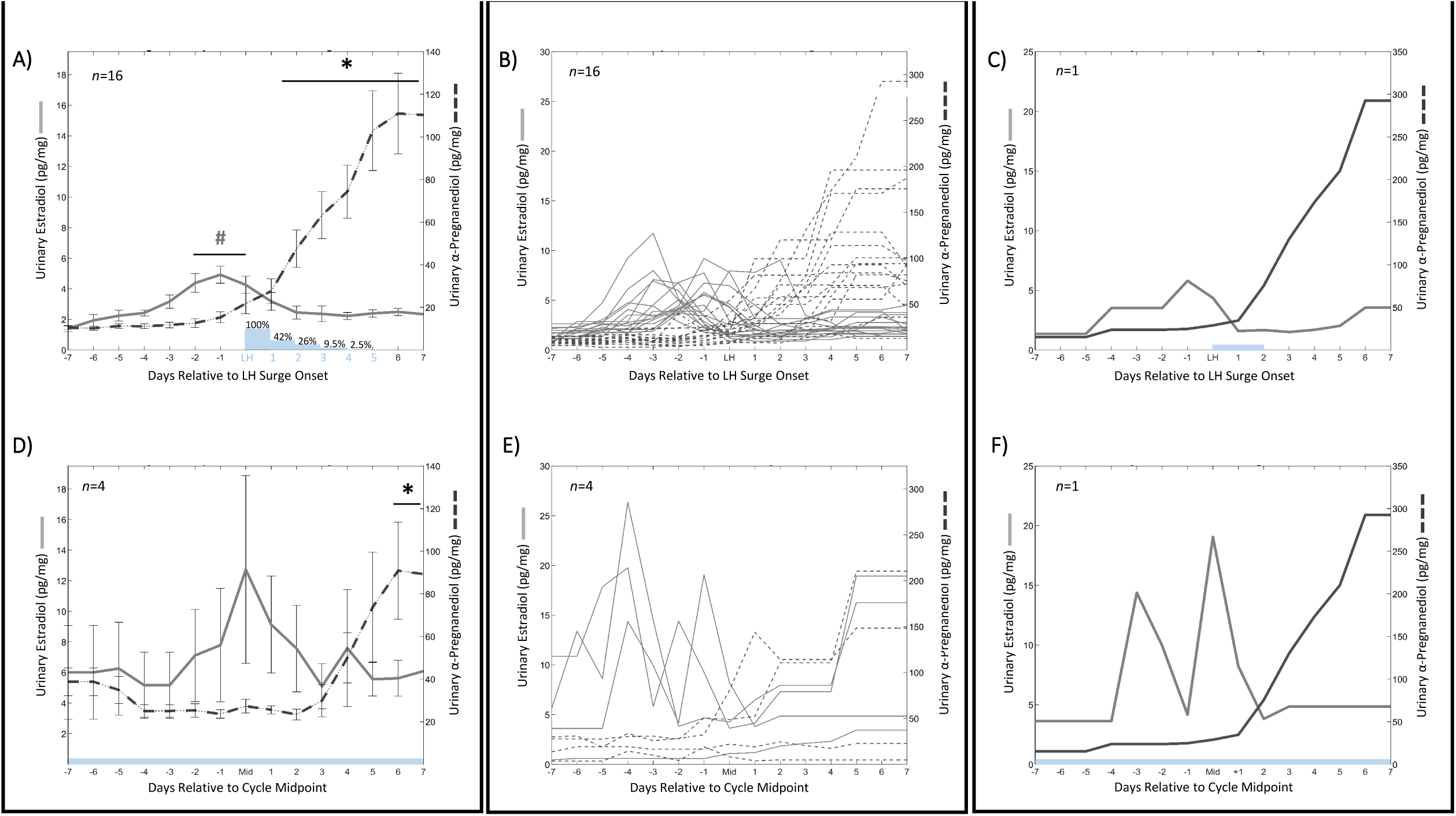
Ovulatory & Perimenopausal Estradiol and a-Pregnanediol. Linear plots of premenopausal (A-C) and perimenopausal (D-F) estradiol and α-pregnanediol. Mean ± standard deviation estradiol (solid) and α-pregnanediol (dashed) concentrations for premenopausal (A) and perimenopausal cycles (D) within one week of LH surge onset (N=16 out of 45 premenopausal cycles, and N=4 out of 10 perimenopausal cycles). # Indicates significantly elevated pre-LH estradiol concentrations (premenopausal *p*=5.5*10^−5^; perimenopausal non-significant *p*=0.391), and * indicates significantly elevated α-pregnanediol after LH surge onset (premenopausal *p*=4.71*10^−31^; perimenopausal, *p*=0.028). Blue bars and text and indicate percent of cycles showing an LH surge a given number of days after onset, beginning on the day marked “LH” (e.g., 26% indicates that 26% of individuals were still surging on the 3^rd^ day after LH surge onset). Representative estradiol (gray) and α-pregnanediol (black) from premenopausal (C) and perimenopausal (F) individuals relative to LH surge onset, and cycle mid-point, respectively.

### Data Analysis

All code and data used in this paper are available at Open Science Framework (https://osf.io/wzf47/). Code was written in Matlab 2019b, Matlab 2020a and Python 3. Wavelet Transform (WT) code was modified from the Jlab toolbox and from Dr. Tanya Leise (Leise, 2013). Briefly, data were imported from the Open Humans framework to Python 3, where HR, HRV (RMSSD) and DBT data were extracted. Data were cleaned in Matlab, with any data points outside +/- 4 standard deviations set to the median value of the prior hour, and any points showing near instantaneous change, as defined by a local abs(derivative) > 10^5^ as an arbitrary cutoff, also set to the median value of the previous hour.

Wavelet transformation (WT) was used to assess the structure of ultradian rhythms of DBT, HR, and HRV (RMSSD) and circadian rhythms in DBT. As DBT shows high plateaus during the sleeping period, and URs during the day, DBT analyses here were used on data collected during the waking hours (see **Supplemental Figure 2**). Conversely, as indicated previously, because the Oura© ring only collects HR and HRV (RMSSD) during sleep, wavelet analyses were restricted to the sleeping window for these metrics. In either case, the excerpted data were compiled from all days of the cycle resulting in one continuous signal representing all days (DBT) or all nights (HR, HRV (RMSSD)). In contrast to Fourier transforms that transform a signal into frequency space without temporal position (i.e., using sine wave components with infinite length), wavelets are constructed with amplitude diminishing to 0 in both directions from center. This property permits frequency strength calculation at a given position. Wavelets can assume many functions (e.g., Mexican hat, square wave, Morse); the present analyses use a Morse wavelet with a low number of oscillations (defined by *β* and *γ*), analogous to wavelets used in previous circadian applications (Leise, 2013). Morse Wavelet parameters of *β* = 5 and *γ* = 3 describe the frequencies of the two waves superimposed to create the wavelet; Additional values of *β* (3–8) and *γ* (2–5) did not alter the findings (data not shown) (Lilly & Olhede, 2012). This low number of oscillations enhances detection of contrast and transitions. The band of the wavelet matrix corresponding to 2-5 h rhythms were averaged in order to create a linear representation of UR WT power over time. This band corresponded with the daily ultradian peak power observed in URs across physiological systems (Brandenberger *et al*., 1987; Grant *et al*., 2018; Goh *et al*., 2019). Potential changes to circadian power of DBT (average power per minute within the 23-25 h band) were additionally assessed prior to extracting days for ultradian-only analyses, but no significant changes across the cycle were detected (see **Supplemental Figure 3**). Because WTs exhibit artifacts at the edges of the data being transformed, only the WT of the second through the second to last days of data were analyzed further. To enable comparisons across cycles of different durations, premenopausal cycles were displayed from LH surge onset minus 7 days to LH onset plus 7 days. As perimenopausal individuals had tonically high LH, and therefore no surge onset to which all individuals could be aligned, each cycle’s midpoint was chosen for alignment.

**Figure 2.**
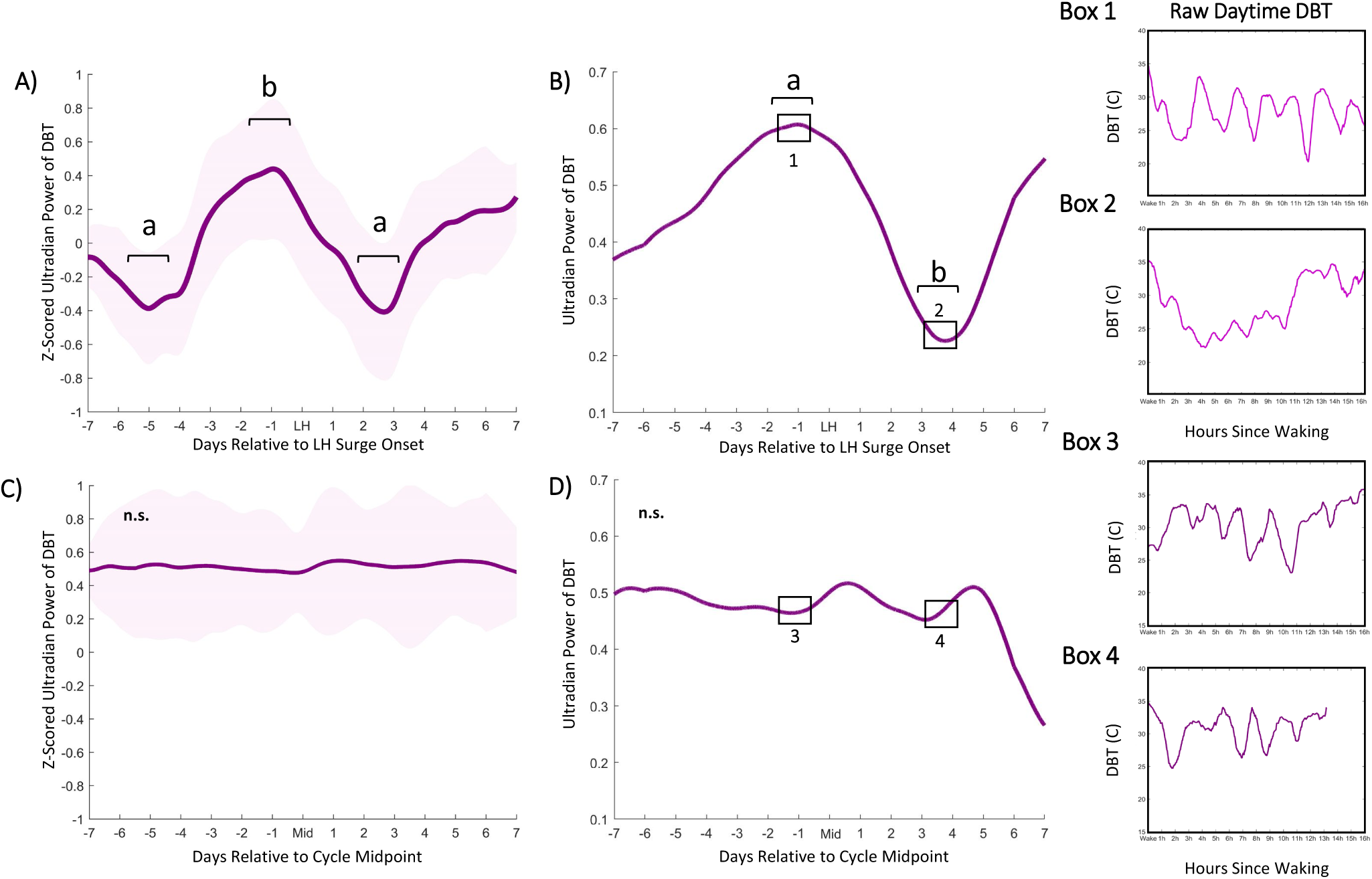
Ultradian Power of DBT Anticipates LH Surge Onset. Mean DBT ultradian power (z-scored)±standard deviation for premenopausal cycles (A) within one week of LH surge onset and perimenopausal cycles (C) within one week of mid cycle. DBT UR power peaks exhibits an inflection point 5.82 (± 1.82) days prior to LH onset, a peak an average of 2.58 (±1.89) before LH onset on average and a subsequent trough an average of 2.6 (±1.02) days after surge onset (χ^2^=5.66, *p*=0.0174). Perimenopausal UR power shows no conserved peaks and troughs (χ^2^= 0.37, *p*=0.5354, for same comparisons). Representative individual example of raw DBT ultradian power within one week of LH surge onset in premenopausal (B) and within one week of mid cycle in perimenopausal (D) cycles. Black squares in (B) and (D) correspond to Boxes 1 & 2 and Boxes 3 & 4, respectively. Boxes show linear waking DBT from which ultradian power in B and D were generated; these days were selected to visually illustrate days of relatively high and low ultradian power in premenopausal cycles, and the same two days in perimenopausal cycles.

**Figure 3.**
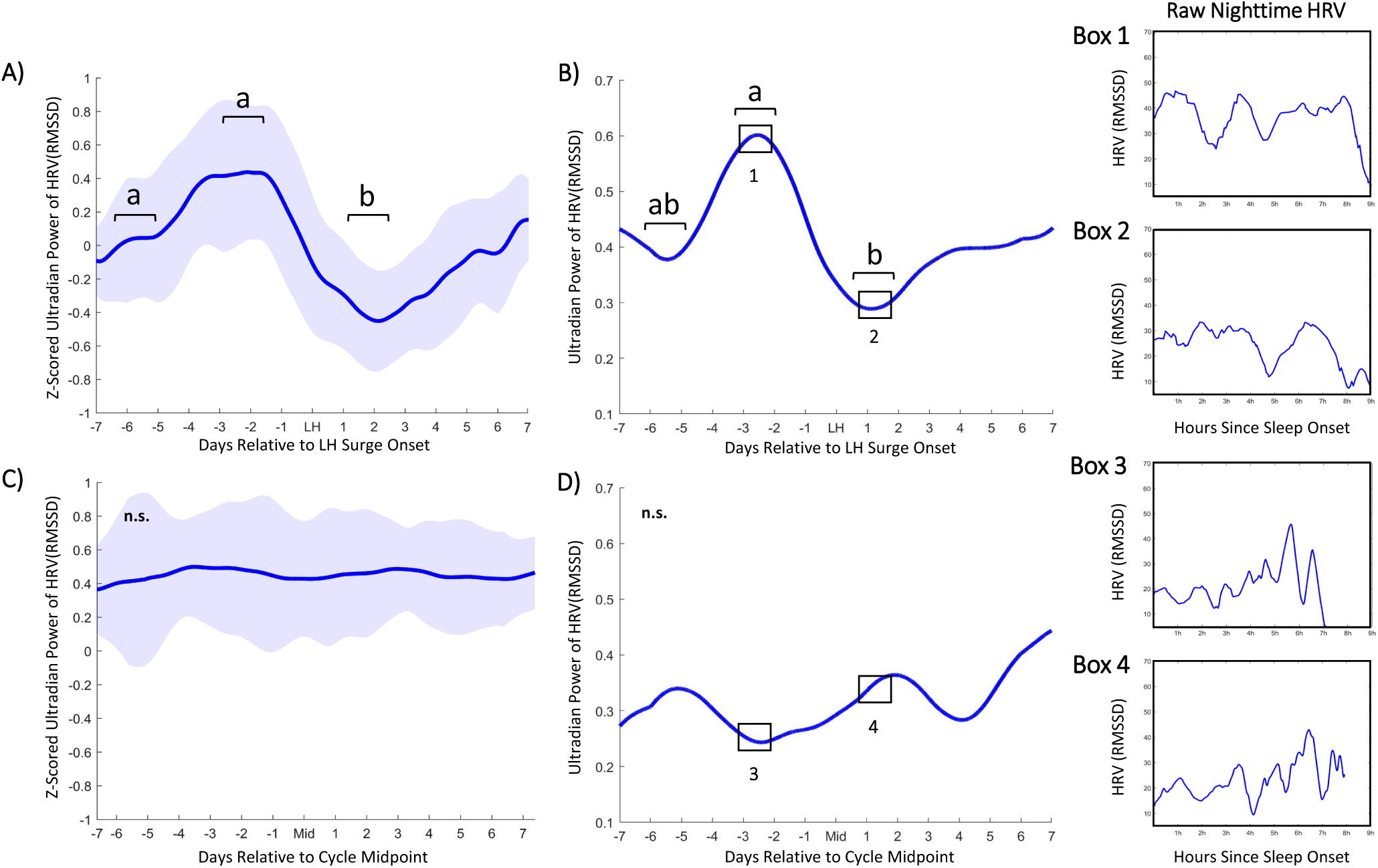
Ultradian Power of HRV (RMSSD) Anticipates LH Surge Onset. Mean HRV (RMSSD) ultradian power (z-scored)±standard deviation for ovulatory cycles (A) within one week of LH surge onset and perimenopausal cycles (C) within one week of mid cycle. Ultradian HRV (RMSSD) power inflects an average of 5.82 (±1.53) nights prior to LH surge onset, exhibits a subsequent peak an average of 2.58 (±1.89) days prior to the surge onset and a trough an average of 2.11 (±1.27) days after surge onset. (χ^2^= 4.91, *p*=0.034,). These stereotyped changes are not present in perimenopausal cycles (χ^2^= 0.4797, *p*=0.57). Representative individual example of HRV (RMSSD) ultradian power within one week of LH surge onset in premenopausal (B) and perimenopausal (D) cycles. Black boxes in (B) and (D) correspond to Boxes 1 & 2 and Boxes 3 & 4, respectively. Boxes show linear sleeping HRV (RMSSD) signal from which B and D were generated, illustrating days of relatively high and low ultradian power in ovulatory cycles, and the same two days in perimenopausal cycles.

### LH Surge Anticipation Features

Wavelet power in the 2-5 h band was calculated as described above. Data were smoothed using a two-day moving average using the Matlab function “movmean”. The Matlab function “findpeaks” was used to identify peaks as points at which either adjacent point had a lower UR power. This function was run on the negative of the signal to identify troughs. Points at which the derivative of the signal crossed zero, indicating a change in direction of UR power (i.e., either increasing to decreasing or vice versa), were found using the matlab function “diff”. The first time the derivative crossed zero in the cycle (i.e., the first inflection point), excluding the first five days of the cycle, during which LH is very unlikely to rise, was marked as the presence of the first feature for either HRV (RMSSD), DBT, or HR. Following this inflection point, the next peak identified by “findpeaks” was marked as the second feature. These methods of identifying peaks, troughs and direction changes were used to ensure the diff function was identifying all visually identified peaks.

### Statistical Analyses

Descriptive values are reported as means ± daily standard deviations (SD) unless otherwise stated. For statistical comparisons of average ultradian power in premenopausal and perimenopausal cycles, Kruskal Wallis (KW) tests were used instead of ANOVAS to avoid assumptions of normality for any distribution to assess the trend in average UR power leading up to the surge as compared to after the surge. For KW tests, *χ* ^2^ and *p* values are listed in the text. One-way repeated measures analysis of variance (rmANOVA) tests were used to compare peak average E2 to other days surrounding the surge, and baseline αPg and βPg (7 days prior to the surge) to other days surrounding the surge. For rmANOVAs, *p* values are listed in the text. Because the dominant trend was an inflection point in UR power followed by a peak, slopes of individual signals were compared rather than raw values at each timepoint. The same tests were applied to individuals, in addition to tests for significance of raw power differences on peak and trough days, using 25 minutes centered on peaks and troughs, respectively. To avoid multiple comparisons and chance of a type I error, differences between individually-determined peak and trough values of UR WT power found using “findpeaks” were assessed using a KW test, such that each cycle contributed only 1 peak value and 1 trough value (N=45 data points per group). Figures were formatted in Microsoft PowerPoint 2019 (Microsoft Inc., Redmond, WA) and Adobe Photoshop CS8 (Adobe Inc, San Jose, CA).

## Results

### Demographics

Findings are reported for individuals with premenopausal cycles (*n*=20, *n*=45 cycles) or perimenopausal cycles as (*n*=5, *n*=10 cycles) as defined above. Individuals who became pregnant (*n*=3) during the study were excluded from the analyses. All premenopausal participants experienced menses, 1 or more supra-threshold LH readings, and a subsequent, sustained rise in temperature deviation during all cycles. See **Table 1** for participant age, ethnicity, cycle length, LH surge length, LH surge onset timing, LH surge onset relative to estradiol peak, and percent of individuals with regular cycles. Some variability was observed in the day of LH surge onset relative to day of estradiol peak(s), as previously reported (Direito *et al*., 2013).

### Premenopausal and Perimenopausal Estradiol, Luteinizing Hormone and Progesterone Metabolites

Participants monitored LH for all 55 cycles, whereas daily urine samples were collected by 20 women (*n*=16 premenopausal, *n*=4 perimenopausal) for the measurement of estradiol, αPg and βPg. Estradiol, *α*Pg and βPg were collected to confirm that hormone concentrations were within healthy ranges for pre-menopausal women and that LH surges were followed by a rise in progesterone metabolites. For all 16 cycles, estradiol, αPg and βPg were within normal ranges, with a pre-LH rise in E2 (2 days prior to LH onset through LH onset day were significantly greater than all other days, *p <*0.01 in all cases). Likewise, LH surge onset was concomitant with a significant rise in αPg (*p* < 0.05 on LH onset, and <0.01 6 days after LH onset and thereafter) (**Figure 1 A-C**) and βPg (*p <* 0.005 on LH onset, and <0.001 3 days after LH onset; data not shown for βPg). These hormonal changes were associated with a rise in temperature deviation above zero, and an elevation of breathing rate at LH surge onset (**Supplemental Figure 1E-F**). Consistent with previous findings (Direito *et al*., 2013), LH surge length was variable, with 42% of individuals exhibiting supra-threshold LH concentrations 2 days following surge onset, falling to 26% of individuals 3 days after surge onset (**Figure 1A**). LH was tonically supra-threshold in perimenopausal women (*n*=10 cycles, **Figure 1D**). Perimenopausal individuals exhibited a significant increase in αPg and βPg only 6 days after midcycle (*p*<0.05; data not shown for βPg), and a trend toward elevation of E2 prior to mid cycle (*p=*0.176).

### Ultradian Power of DBT, HRV and LH Surge Onset: Premenopausal and Perimenopausal Cycles

Ultradian (1-4 h) power of daytime DBT exhibited a stereotyped pattern preceding LH surge onset in premenopausal (**Figure 2 A,B**), but not perimenopausal (**Figure 2 C,D**), cycles. Ultradian DBT power exhibited an inflection point an average of 5.82 (±1.82) days prior to LH surge onset, a subsequent peak an average of 2.58 (±1.89) days prior to the surge onset, and a second trough in UR power occurred an average of 2.06 (±1.02) days after surge onset (χ^2^=5.66, p=0.0174,). These stereotyped changes were not present in perimenopausal cycles (χ^2^= 0.37, p=0.5354, for the same comparisons) (**Figure 2C**).

Ultradian (1-4 h) power of sleeping HRV (RMSSD) exhibited a stereotyped fluctuation preceding LH surge onset in premenopausal (**Figure 3A,B**), but not perimenopausal (**Figure 3C,D**) cycles. Ultradian HRV (RMSSD) power showed an inflection point an average of 5.82 (±1.53) nights prior to LH surge onset, a subsequent peak an average of 2.58 (±1.89) days prior to the surge onset and a trough an average of 2.11 (±1.27) days after surge onset. (χ^2^= 4.91, p=0.034). These stereotyped changes were not present in perimenopausal cycles (χ^2^= 0.4797, p=0.57) (**Figure 3C**). Ultradian power of HR and circadian power of DBT did not show a significant pattern of change preceding the LH surge (χ^2^= 0.3 and 1.12, p=0.581 and 0.2899), nor mid cycle in perimenopausal individuals (χ^2^= 0.02 and 1.65, p= 0.8798 and 0.1984, respectively) (See **Supplemental Figure 2 and 3**). No significant trends were observed in sleep metrics captured once per night (See **Supplemental Figure 1**). Linear averages of nightly HR and HRV, and continuous DBT did not yield consistent patterns of change relative to surge onset or perimenopausal midcycle (**Supplemental Figures 4-6**).

**Figure 4.**
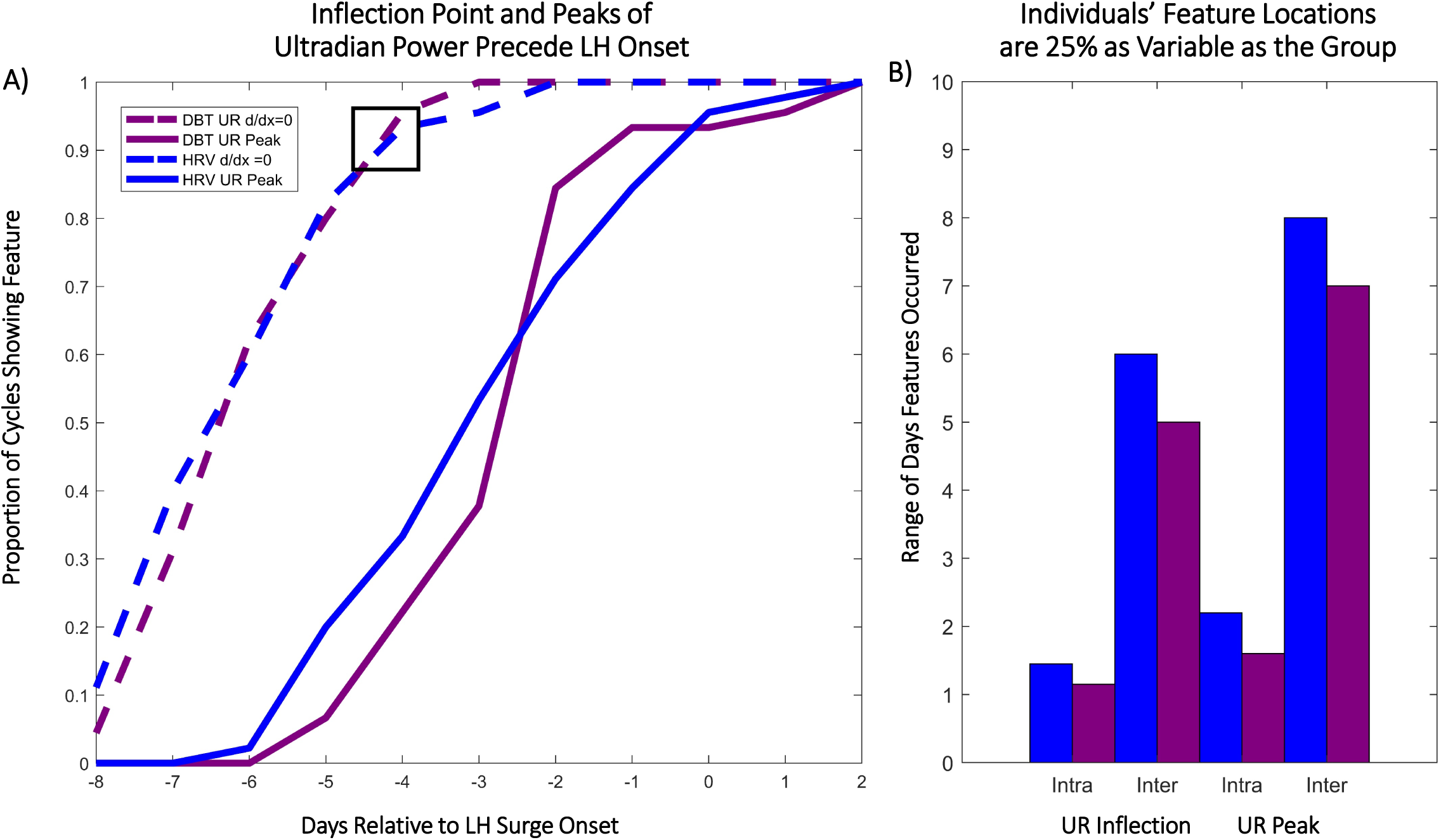
Inflection Points and Peaks of Ultradian Power Anticipate the LH Surge Within and Across Individuals. A) Cumulative histogram indicates the proportion of cycles showing an inflection point on a given day relative to LH surge onset (blue=HRV (RMSSD), maroon=DBT, dashed = inflection point, solid= subsequent peak). Box indicates that on day LH – 4, ∼90% of individuals had shown the HRV and DBT first inflection point. B) Intra-vs. Inter Individual range of days over which inflection points (“UR Inflection”) and subsequent peaks (“UR Peak”) of ultradian DBT and HRV power occurred. The range intra-individual range (2-3 cycles per individual) is 25% the size of the inter-individual range (45 total cycles).

### Inflection Point and Subsequent Peak of DBT and HRV Ultradian Power Anticipate LH Surge Onset

In premenopausal women, the first inflection point of DBT and HRV (RMSSD) UR power occurred between −8 and −2 days prior to surge onset, whereas the subsequent peak in UR power for both metrics occurred between −6 days before to 2 days after LH surge onset (**Figure 4a**). 85% of cycles exhibited the first inflection point by 4 days prior to the surge, with 100% showing this inflection point by 2 days prior to the surge. The peak of UR power occurred at least 1 day prior to the surge in 82% of cycles. Together, these inflection points and subsequent peaks in UR power of HRV (RMSSD) and DBT uniquely anticipated the LH surge days before its onset (See Discussion for potential relevance to fertile window).

## Discussion

The present findings reveal stereotyped fluctuations in DBT and HRV (RMSSD) UR power that anticipate 100% of LH surge onsets, a key component of female health and fertility. In contrast, changes in DBT circadian rhythm power were not predictive of the LH surge, suggesting that URs are uniquely coupled to the pre-ovulatory time of the menstrual cycle. Likewise, discrete, nightly behavioral and physiological measures did not anticipate the surge, suggesting that continuous measures of physiological output provide signals more amenable to LH surge anticipation. Finally, these features could not be identified in perimenopausal cycles with respect to mid cycle. These findings point to URs as oscillations that are coupled to menstrual cycle physiology and that have the potential to contribute to the development of tools for estimating the female fertile window.

Although the underlying physiological mechanisms that lead to systematic changes in UR power in DBT require further investigation, much is known about general changes in body temperature across the menstrual cycle. Estrogens lower, and progestins raise, body temperature (Buxton & Atkinson, 1948; Williams *et al*., 2010). Accordingly, body temperature reaches a minimum, with minimum core circadian amplitude, during the late follicular phase and rises in the core, mouth, and skin following ovulation (Coyne *et al*., 2000). Body temperature also broadly reflects metabolic rate, which is elevated in the late follicular and luteal phases (Solomon *et al*., 1982). In mice, the structure of core temperature URs allows for the detection of female reproductive state, with a high plateau of temperature during the active phase indicative of the LH surge and ovulation (Sanchez-Alavez *et al*., 2011; Smarr *et al*., 2017). Most studies to date have focused on core temperature, measured via an ingestible device that travels through the GI tract (Coyne *et al*., 2000), intravaginal or rectal sensor (Shechter & Boivin, 2010), or oral thermometer (Kräuchi *et al*., 2014). However, ultradian, circadian and ovulatory rhythms in temperature are readily observed at the periphery, providing several advantages: 1) DBT has higher amplitude fluctuations than core body temperature, making URs/CRs easier to detect (Szymusiak, 2018), 2) Changes in DBT correlate with sleep stage (Henane *et al*., 1977), and 3) DBT is in circadian antiphase to core temperature, but shows the same general trend across the menstrual cycle, suggesting comparable reliability (Szymusiak, 2018).

Whereas body temperature is the most commonly used non-hormonal output in menstrual cycle tracking, previous studies have found that HRV also changes by cycle phase and may therefore be a candidate for surge anticipation (Schmalenberger *et al*., 2019). Parasympathetic input to the heart dominates during the follicular phase, lowering resting heart rate and elevating HRV (RMSSD) (Sato *et al*., 1995). Sympathetic input to the heart dominates during the luteal phase, elevating heart rate and depressing HRV (RMSSD) (Sato *et al*., 1995; Brar *et al*., 2015). Consequently, HRV (RMSSD) varies ∼ 10 ms from the follicular to the luteal phase (Brar *et al*., 2015), with a marked decrease 14 days before the end of the cycle (Visrutha *et al*., 2012). These fluctuations may be more difficult to detect during a short daytime recording window, and are impacted by daytime activities, making the sleeping period an ideal window over which to look for unmasked features (Leicht *et al*., 2003). Natural negative controls illustrating reproductive and metabolic influence on HRV pattern are 1) that LH pulsatility is disrupted in obese and diabetic women (Jain *et al*., 2007; van Leckwyck *et al*., 2016), and 2) mid cycle and luteal fluctuations in HRV are absent in polycystic ovarian syndrome (PCOS), the leading cause of female infertility (Yildirir *et al*., 2006; Uckuyu *et al*., 2013). In the present study, sleeping HRV (RMSSD) UR power rose in the late follicular phase, peaked near the LH surge, and dropped sharply before rising into the early luteal phase. Together, signal processing of DBT and HRV could yield actionable information for individuals and clinicians wishing to estimate the “fertile window”. However, there are several challenges inherent to accurately defining the female fertile window.

The fertile window (the time during which a woman may become pregnant) depends upon many factors, including 1) the time of the LH surge, 2) the subsequent time of the release of the ovum or ovulation, 3) the presence of a viable corpus luteum releasing adequate progesterone (Soules *et al*., 1989), 4) the duration of time sperm can survive in the female body, which is dependent both on sufficient number and quality of sperm and on the appropriate vaginal environment (e.g., pH) (Stanford, 2015; Lessey & Young, 2019), and 5) quality of the uterine environment. Most investigations report the highest probability of fertility as the 5 days preceding ultrasound-determined day of ovulation (USDO) (Faust *et al*., 2019), but actual days on which an individual may become pregnant are much more variable, with pregnancy occurring 11 days prior to ovulation to 5 days after ovulation (Direito *et al*., 2013).

Some of the reported variability in the fertile window likely results from discrepancies in language used to describe both human ovulation and the fertile window itself (Setton *et al*., 2016). Despite their namesake, home “ovulation tests” that identify supra-threshold LH concentrations, for example, do not measure ovulation, which may occur many days after surge onset and occasionally a few days *before* LH surge onset (Direito *et al*., 2013). Despite this variability, the fertile window is often treated as predictable, with definitions including the 5-6 days prior to the LH surge as a proxy for ovulation (Su *et al*., 2017), the first day of slippery clear cervical mucus through LH onset (Keulers *et al*., 2007), the total days of slippery clear cervical mucus (Ecochard *et al*., 2015), day 10-17 of the cycle (Wilcox *et al*., 2000), and retrospective measures of salivary ferning (Su *et al*., 2017), basal body temperature (Moghissi, 1976; Quagliarello & Arny, 1986), and progesterone metabolites^14^. Today, many online and app-based ovulation prediction algorithms are validated using day of cycle or LH data alone, in the absence of hormone measures or USDO (Setton *et al*., 2016; Shilaih *et al*., 2017; Freis *et al*., 2018; Faust *et al*., 2019). Additionally, extant data sets regularly report excluding 20-50% of collected data due to cycle irregularities, without determining if given cycles were hormonally aberrant (Renaud *et al*., 1980; Queenan *et al*., 1980; Polan *et al*., 1982; Direito *et al*., 2013). Together, the confounding of the LH surge with the ovulation and the variable criteria used to define the fertile window make it difficult to accurately determine the variance, and contributors to variability, of fertility relative to ovulation.

Open source, non-invasive methods for predicting the LH surge as a marker of likely future ovulation are not currently available (Direito *et al*., 2013), but the present findings indicate that the onset of the LH surge may be anticipated days in advance by automated detection of changes in ultradian power in DBT and HRV (RMSSD). These changes consistently anticipate LH surge onset in women of a variety of ages, contraceptive use history, cycle length, surge timing and duration, pointing to potentially broad applicability for this method of detection. Due to the high demand for accurate methods of fertility assessment, such novel methods carry the responsibility to clearly report the aspects of reproductive physiology that are detected and the methods by which detection is achieved once algorithms are tested on large populations (Setton *et al*., 2016; Epstein *et al*., 2017; Freis *et al*., 2018). Together, these findings may guide further research aimed at understanding how hormones, metabolism and the autonomic nervous system temporally interact; and may aid the development of open-source, non-invasive methods of fertility awareness.

## Acknowledgements

The authors would like to thank the participants in the study for their interest, careful longitudinal self-tracking, and excellent questions and feedback. We also thank Dr. Linda Wilbrecht and Dr. Andrew Ahn for their helpful suggestions on an earlier version of this manuscript.

## Supplemental Figure Legends

**Supplemental Figure 1.**
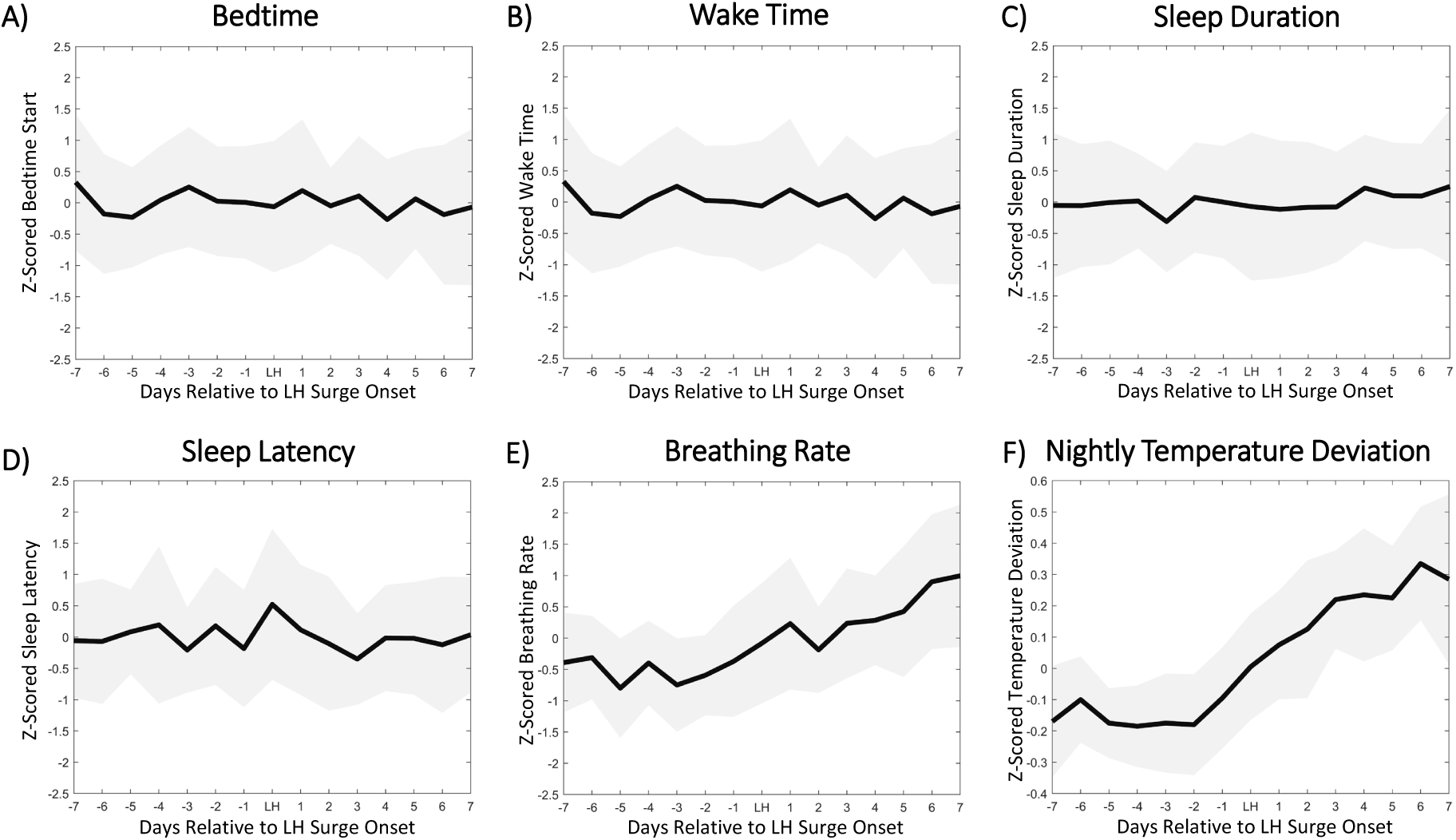
Sleep Timing, Duration, Latency, Breathing Rate and Temperature Deviation do not anticipate the LH Surge. *S*. Bedtime (A), wake time (B), sleep duration (C) and sleep latency (D) show no concerted changes relative to LH surge onset in premenopausal individuals. Breathing rate (E) and nightly temperature deviation (F, see methods) do not anticipate the surge, but exhibit an upward trend following the surge, as previously described (Maijala *et al*., 2019).

**Supplemental Figure 2.**
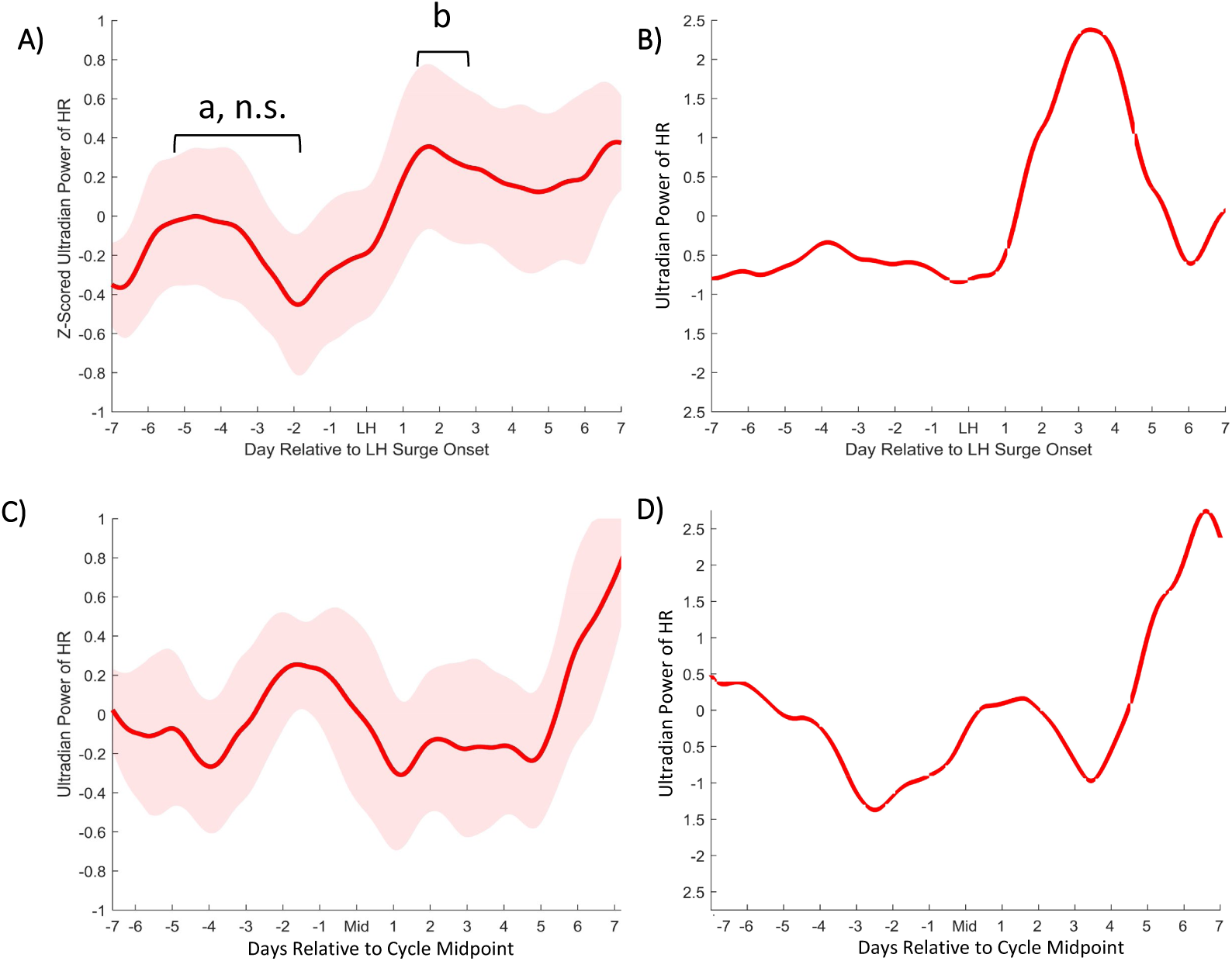
Heart Rate Ultradian Power does Not Anticipate LH Surge Onset. Mean sleeping heart rate ultradian power ± standard deviation for premenopausal cycles (A) within one week of LH surge onset and perimenopausal cycles (C) within one week of mid cycle. HR ultradian fluctuations did not anticipate the LH surge (p>0.05) but exhibited a significant elevation 2-3 days after the surge (χ^2^ =0.3, *p*=0.04). Representative individual examples of HR ultradian power within one week of LH surge onset or mid cycle in premenopausal (B) and perimenopausal (D) cycles, respectively.

**Supplemental Figure 3.**
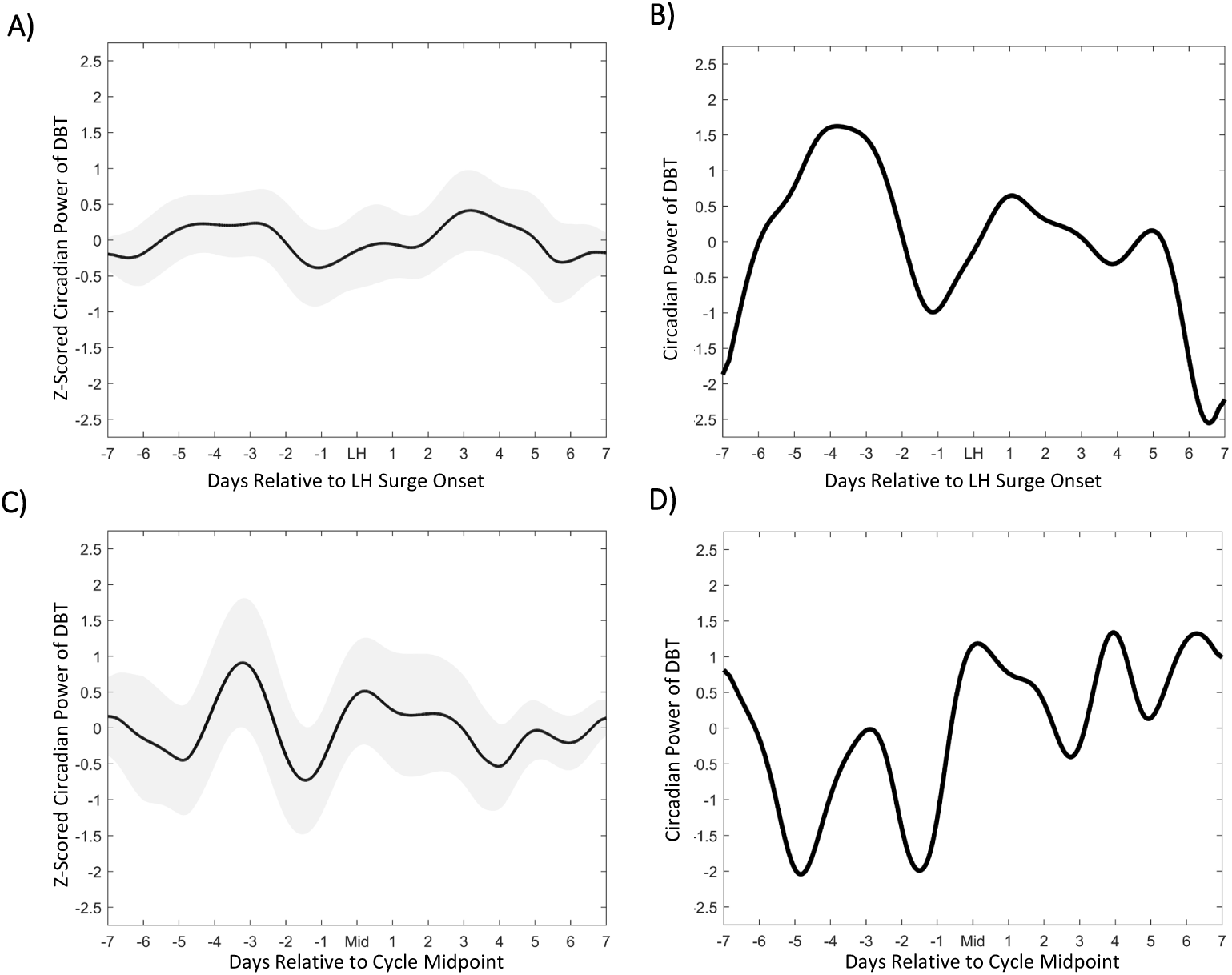
Circadian Power of Body Temperature Does Not Change Stereotypically Around LH Surge Onset. Mean DBT circadian power ± standard deviation for premenopausal cycles (A) within one week of LH surge onset and perimenopausal cycles (C) within one week of mid cycle did not exhibit significant stereotyped fluctuations relative to LH surge onset or mid cycle. Individuals varied widely (examples B and D).

**Supplemental Figure 4.**
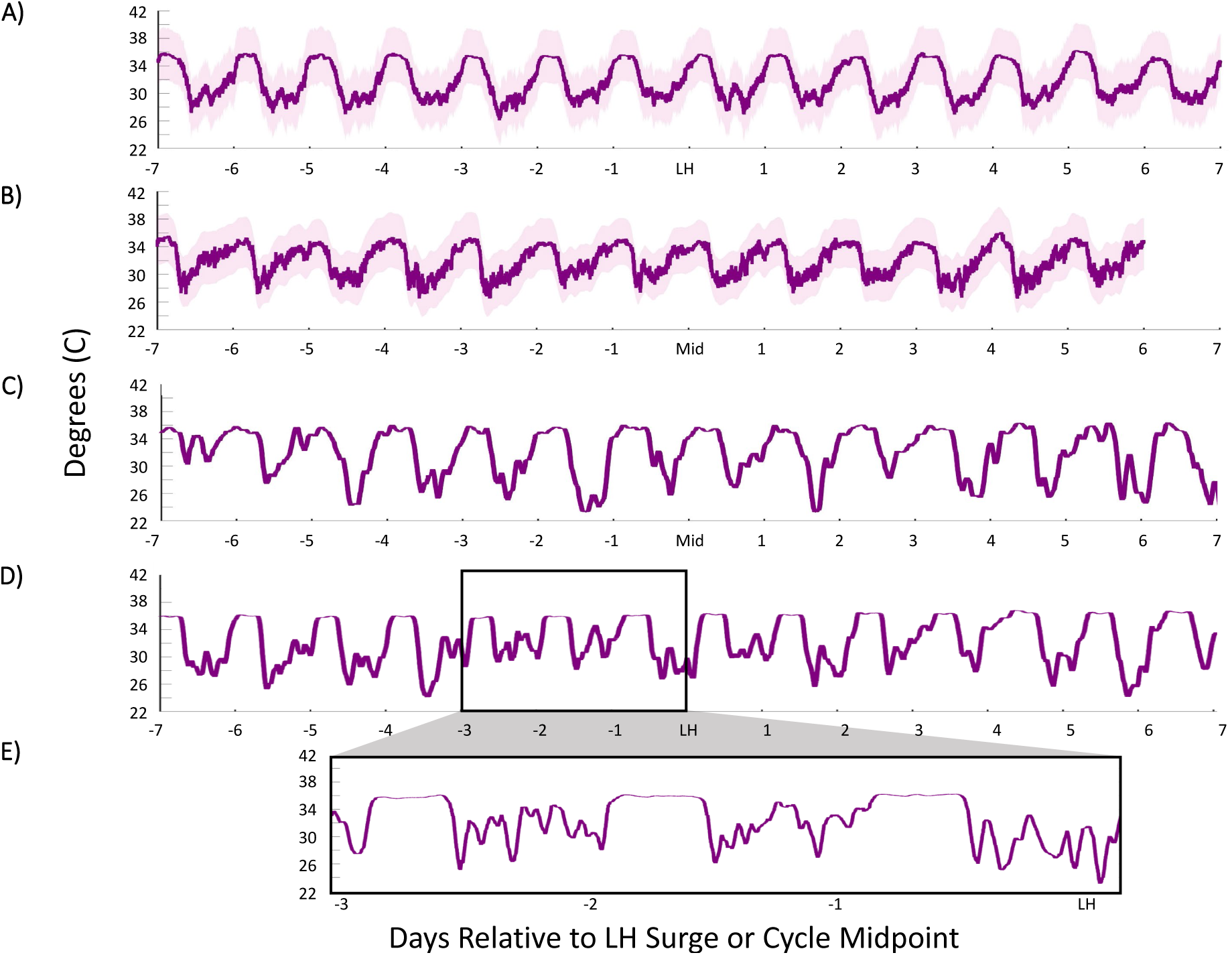
Linear Average of DBT Relative to LH Surge Onset in Premenopausal and Mid Cycle in Perimenopausal Individuals. Mean (A), ± standard deviation (shaded) of linear DBT around the LH surge onset. Perimenopausal mean (B) ± standard deviation (shaded) of linear DBT surrounding mid cycle (note two perimenopausal cycles were very short, with only 6 days after midcycle occurring before next menses), and individual example (C). Individual example of DBT around LH surge onset (D) and zoomed window if this individual from LH-3 days to LH illustrating the presence of high amplitude URs during the day and a relatively high plateau during sleep (E).

**Supplemental Figure 5.**
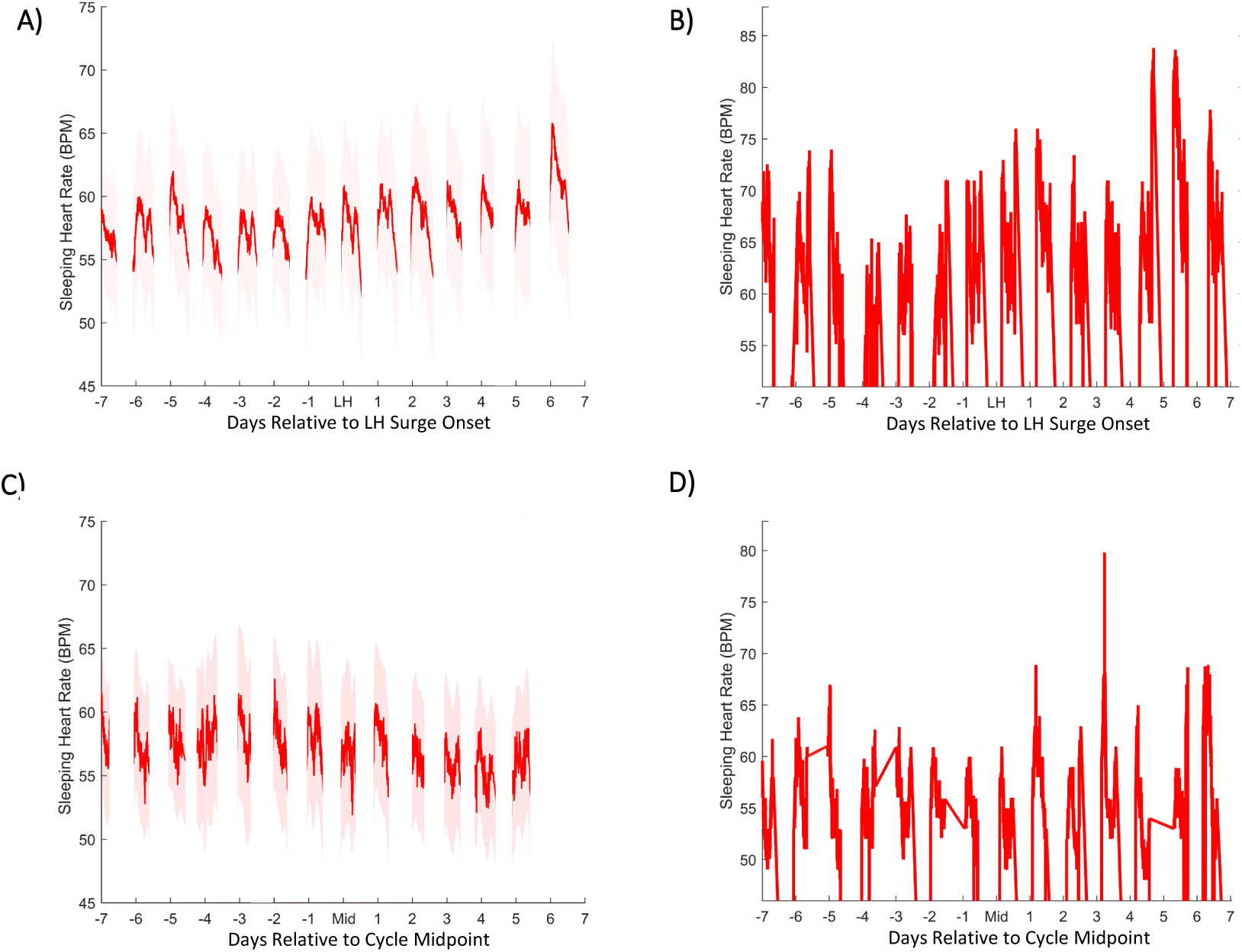
Linear Average of Sleeping HR Relative to LH Surge Onset in Premenopausal and Mid Cycle in Perimenopausal Individuals. Premenopausal average (A), ± standard deviation (shaded) of linear Sleeping HR in surrounding the LH surge onset, and individual example (B). Perimenopausal average (C) ± standard deviation (shaded) of linear HR surrounding mid cycle, and individual example (D).

**Supplemental Figure 6.**
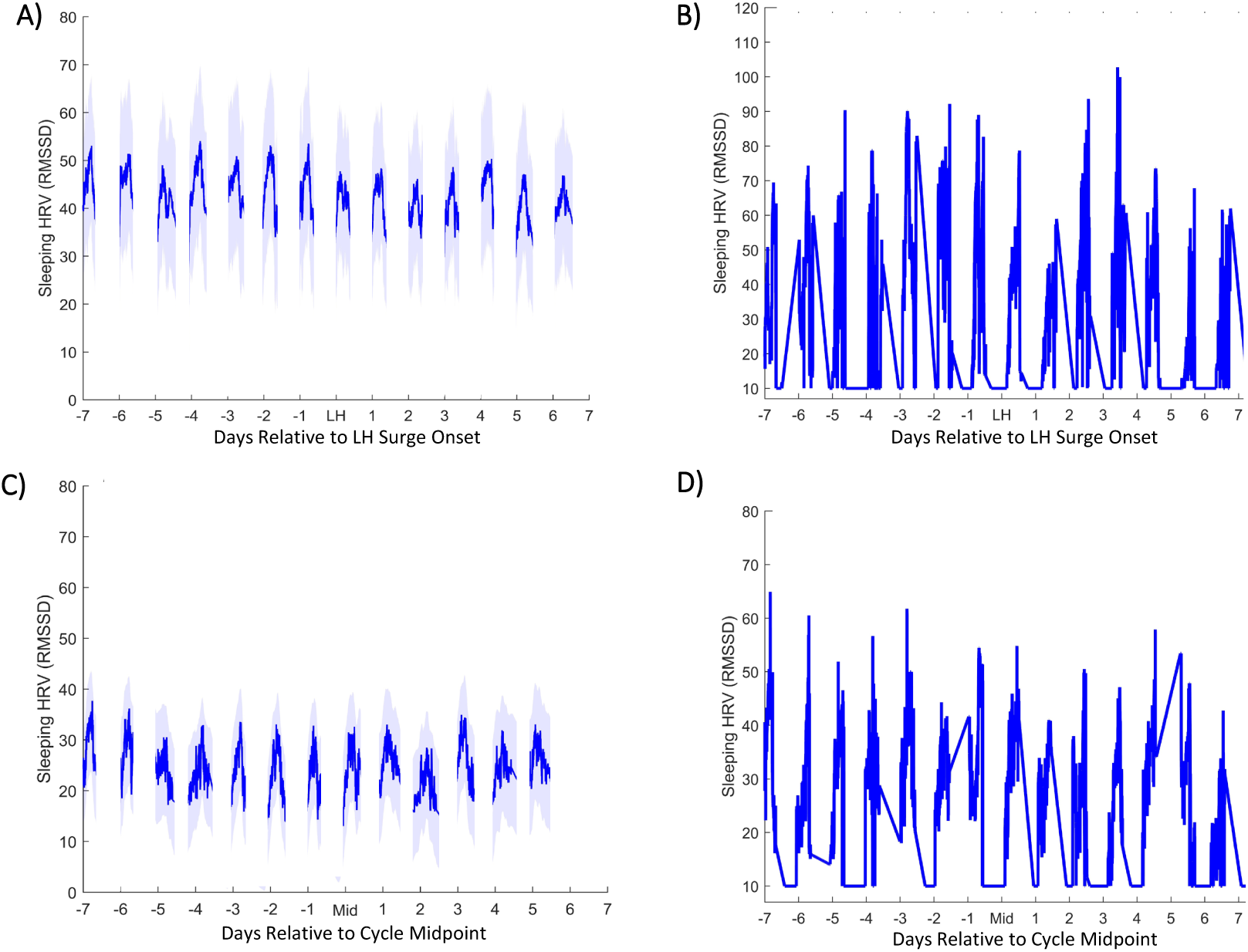
Linear Average of Sleeping HRV (RMSSD) Relative to LH Surge Onset in Premenopausal and Mid Cycle in Perimenopausal Individuals. Premenopausal average (A), ± standard deviation (shaded) of linear Sleeping HRV in surrounding the LH surge onset, and individual example (B). Perimenopausal average (C) ± standard deviation (shaded) of linear HRV surrounding mid cycle, and individual example (D).

## LH Surge Anticipation Figures

**Table 1.**

| Factor | Premenopausal | Perimenopausal |
| --- | --- | --- |
| Number of Participants | 20 | 6 |
| Number of Cycles | 45 | 10 |
| Age Range, Mean (STDEV) *years* | 21-38, 32 (4) | 48-60, 55 (5) |
| Ethnicity (%) | Caucasian: 94; African:<br>American 6 | Caucasian: 100% |
| Cycle Length | 25-36, 27.78 (4.16) | 22-50, 28.7 (8.87) |
| LH Surge Length Range, Mean<br>(STDEV) *days* | 1-5,1.95 (1.2) | N/A; LH Tonically High |
| LH Surge Onset Day | 10-29,15.75 (3.4) | N/A; LH Tonically High |
| LH Surge Onset Relative to E2<br>Peak in *days* | 0-4,1.67,(1.38) | N/A ; LH Tonically High |
| LH Surge Day Relative to<br>Progesterone Rise *day* (n=21) | 0-5,1.14,(1.95) | N/A; LH Tonically High |
| Regular Cyclers (%) | 88 | 0 |

